# *Mycobacterium tuberculosis* response to cholesterol is integrated with environmental pH and potassium levels via a lipid utilization regulator

**DOI:** 10.1101/2023.08.22.554309

**Authors:** Yue Chen, Nathan J. MacGilvary, Shumin Tan

## Abstract

How bacterial response to environmental cues and nutritional sources may be integrated in enabling host colonization is poorly understood. Exploiting a reporter-based screen, we discovered that overexpression of *Mycobacterium tuberculosis* (Mtb) lipid utilization regulators altered Mtb acidic pH response dampening by low environmental potassium (K^+^). Transcriptional analyses unveiled amplification of Mtb response to acidic pH in the presence of cholesterol, a major carbon source for Mtb during infection, and vice versa. Strikingly, deletion of the putative lipid regulator *mce3R* resulted in loss of augmentation of (i) cholesterol response at acidic pH, and (ii) low [K^+^] response by cholesterol, with minimal effect on Mtb response to each signal individually. Finally, the Δ*mce3R* mutant was attenuated for colonization in a murine model that recapitulates lesions with lipid-rich foamy macrophages. These findings reveal critical coordination between bacterial response to environmental and nutritional cues, and establish Mce3R as a crucial integrator of this process.

## INTRODUCTION

Bacterial colonization of its host requires adaptation of the bacteria to their local environment, which includes ionic signals such as protons (H^+^, pH levels), chloride (Cl^−^), and potassium (K^+^)^1–3^. *Mycobacterium tuberculosis* (Mtb) is the causative agent of tuberculosis, a disease that continues to be a leading cause of death from infectious diseases worldwide^4^. The importance of appropriate sensing and response to the local environment for Mtb is underlined by the substantial attenuation in colonization ability of a mutant deleted for *phoPR*^5,6^, a two-component system essential in the response of Mtb to acidic pH that also plays a significant role in its response to Cl^−^^1,2^. Indeed, the attenuation of a Δ*phoPR* mutant for host colonization is so marked that it has been pursued as a possible vaccine candidate^7,8^. Environmental cues further change in time and space during infection, and we have previously shown that spatial variations in local pH and Cl^−^ levels exist even within a single lesion, and correlate to Mtb replication status^9^.

Critically, bacteria are exposed during infection not just to a single signal at a time, but concurrently to multiple cues in their local environment. In the case of Mtb, we have shown that the bacteria have a synergistic transcriptional response to the simultaneous presence of acidic pH and high Cl^−^ concentration ([Cl^−^]), two signals experienced concurrently within a maturing macrophage phagosome^2^. There is further a relationship between the K^+^ homeostasis of Mtb and its response to acidic pH/high [Cl^−^], with deletion of the CeoBC K^+^ uptake system impairing the response of Mtb to these two environmental cues and negatively affecting the ability of Mtb to colonize its host^3^. Besides PhoPR regulating Mtb response to both acidic pH and high [Cl^−^]^2^, other examples of coordinated regulation in signal response are exemplified by additional key two-component systems such as DosRS(T) (nitric oxide stress and hypoxia) and PrrAB (acidic pH, high [Cl^−^], nitric oxide stress, hypoxia)^10,11^. Of note, PrrA is essential in Mtb^11,12^, while mutants in DosRS(T) show significant attenuation in several animal models of Mtb infection^13–15^. These data emphasize the existence of interconnections between environmental signals, and the importance of coordination in response to these signals for successful host colonization by Mtb.

Alongside environmental stimuli, the availability of nutrients also plays a crucial role in bacterial adaptation to its host. Particularly for Mtb, cholesterol has been shown to be a major carbon source during infection, with Mtb mutants deficient in their ability to utilize cholesterol attenuated for host colonization^16,17^. Strikingly, Mtb exhibits improved growth in acidic pH media when cholesterol or fatty acids are provided as carbon sources^18,19^. Further, the transcriptional response of Mtb to Cl^−^ was increased in the simultaneous presence of cholesterol^20^. In addition, two studies have found that iron limitation increases expression of the cholesterol utilization genes in Mtb^21,22^. Together, these data suggest the presence of intrinsic links between Mtb response to environmental (ionic) cues and its metabolism. However, the nature of these relationships and the regulatory factors that underlie them remain largely unexplored and unknown.

Here, building on our previous finding that disruption of Mtb K^+^ homeostasis decreases the bacterial response to acidic pH and high [Cl^−^]^3^, we first discover that limiting environmental K^+^ levels also dampen Mtb response to acidic pH. Exploitation of a pH reporter Mtb-based transcription factor overexpression screen then unexpectedly revealed a connection between Mtb lipid metabolism regulation with control of the bacterial pH response in the presence of limiting [K^+^]. Focusing on the putative lipid utilization regulator Mce3R, overexpression of *mce3R* partially rescued, while *mce3R* deletion conversely further repressed, the acidic pH response of Mtb in the presence of low external [K^+^]. The presence of cholesterol was found to increase Mtb transcriptional response to low [K^+^], with global transcriptional analyses demonstrating that Mtb response to acidic pH was also augmented in the simultaneous presence of cholesterol and vice versa. Strikingly, deletion of *mce3R* resulted in loss of augmentation of the cholesterol response in the presence of acidic pH, while having minimal effect on Mtb transcriptional response to cholesterol or acidic pH alone. Similarly, augmentation of the low [K^+^] response by the simultaneous presence of cholesterol was lost in the Δ*mce3R* Mtb mutant. Lastly, we show that deletion of *mce3R* had significant consequences for Mtb host colonization, with the Δ*mce3R* mutant attenuated for growth in foamy macrophages and colonization of C3HeB/FeJ mice during a six-week infection. Together, our findings reveal the intrinsic interconnections between Mtb response to acidic pH, low [K^+^], and cholesterol, and establish Mce3R as a critical mechanism through which integration of these signals is accomplished in Mtb.

## RESULTS

### Environmental K^+^ levels modulate Mtb transcriptional response to acidic pH

To investigate the impact of environmental potassium (K^+^) levels on the response of Mtb to acidic pH, we exposed our previously established pH-responsive Mtb reporter, *rv2390c’*::GFP^2^, to standard 7H9, pH 7 medium, or 7H9, pH 5.85 medium with sufficient K^+^ (7.6 mM) or a K^+^ concentration limited to 0.1 mM. As shown in Figure 1A, K^+^ limitation strikingly decreased the induction of *rv2390c’*::GFP fluorescence signal observed at acidic pH. This decrease in acidic pH response was further [K^+^]-dependent, with increased dampening of reporter induction with decreasing [K^+^] (Figure 1B). Importantly, this effect of K^+^ limitation on the acidic pH response of Mtb was unidirectional, as the bacterial response to low [K^+^], tested using the K^+^ reporter Mtb strain *kdpF’*::GFP^3^, was not influenced by changes in environmental pH (Figure 1C).

**Figure 1.**
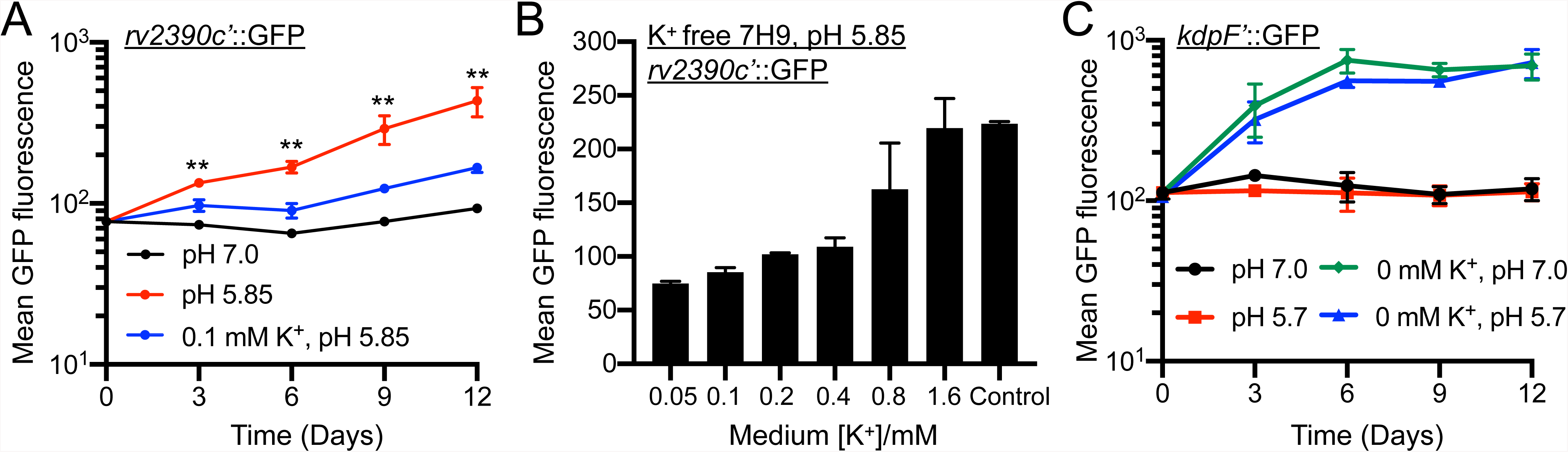
Environmental K^+^ levels modulate Mtb transcriptional response to acidic pH. (A) *rv2390c’*::GFP reporter response to acidic pH is dampened in the presence of low [K^+^]. Mtb(*rv2390c*’::GFP) was grown in 7H9, pH 7.0 or pH 5.85 media, or in K^+^-free 7H9, pH 5.85 medium, supplemented with 0.1 mM K^+^. Samples were taken at indicated time points, fixed, and GFP induction analyzed by flow cytometry. Data are shown as means ± SD from 3 experiments. p-values were obtained with an unpaired t-test with Welch’s correction, and compare the 7H9, pH 5.85 to the 0.1 mM K^+^, pH 5.85 condition. ** p<0.01. (B) Dampening of Mtb response to acidic pH by low K^+^ is concentration dependent. Mtb(*rv2390c*’::GFP) was grown in 7H9, pH 5.85 medium (“control”), or in K^+^-free 7H9, pH 5.85 medium, supplemented with indicated amounts of K^+^. Samples were taken 9 days post-assay start, fixed, and GFP induction analyzed by flow cytometry. Data are shown as means ± SD from 3 experiments. (C) Mtb response to low [K^+^] is not affected by environmental pH. Mtb(*kdpF*’::GFP) was grown in 7H9, pH 7.0 or pH 5.7, or in K^+^-free 7H9, pH 7 or pH 5.7 media. Samples were taken at indicated time points, fixed, and GFP induction analyzed by flow cytometry. Data are shown as means ± SD from 3 experiments.

Together, these assays reveal a previously unknown impact of environmental [K^+^] levels on the response of Mtb to acidic pH.

### Lipid utilization regulators modulate the interplay between environmental [K^+^] and Mtb pH response

To investigate the regulatory mechanism(s) underlying the relationship between environmental [K^+^] and the acidic pH response of Mtb, a screening approach was employed using an arrayed inducible transcription factor overexpression library generated in the background of a pH-responsive *rv2390c’*::luciferase reporter^11^. 7H9, pH 7, 0.05 mM K^+^ or 7H9, pH 5.85, 1.6 mM K^+^ media served as negative and positive control conditions respectively, with an empty vector control included. Two K^+^ limiting concentrations, 0.1 and 0.05 mM, both at pH 5.85, were used as the test conditions. The fold induction of *rv2390c’*::luciferase reporter signal in each test condition compared to the negative control was calculated as relative light units (RLU)/OD_600_ to account for any variations in bacterial growth. The arrayed set-up of the screen removed any concerns of bottleneck effects and enabled immediate identification of hits of interest.

Unexpectedly, a key class of hits identified from the screen were regulators involved in lipid (cholesterol or fatty acid) metabolism (Figure 2A, Table S1). In particular, overexpression of *rv1219c* and *mce3R* partially rescued the dampening of *rv2390c’*::luciferase signal at pH 5.85 with limiting [K^+^] (Figure 2A, Table S1). Conversely, overexpression of *mce1R*, *kstR*, *mce2R*, *rv1129c*, and *rv1816* repressed *rv2390c’*::luciferase induction even more compared to the empty vector control (Figure 2A, Table S1). These phenotypes were first validated by comparing uninduced (ethanol, EtOH) conditions versus induced (200 ng/ml anhydrotetracycline, ATC) conditions for each transcription factor overexpression strain, with outcomes consistent with the screen results (comparison to empty vector) obtained in all cases (Figure 2B). As noted above, these seven transcription factors are known or predicted to be involved in lipid metabolism. Specifically, *mce1R, mce2R*, *rv1219c*, and *rv1816* have previously been shown to be involved in fatty acid metabolism^23–27^, *kstR* is a key regulator of cholesterol β-oxidation^28,29^, and *rv1129c* is involved in propionyl-CoA metabolism^30^. *mce3R* is predicted to be involved in lipid metabolism due to its homology to the other Mce complexes^31–34^.

**Figure 2.**
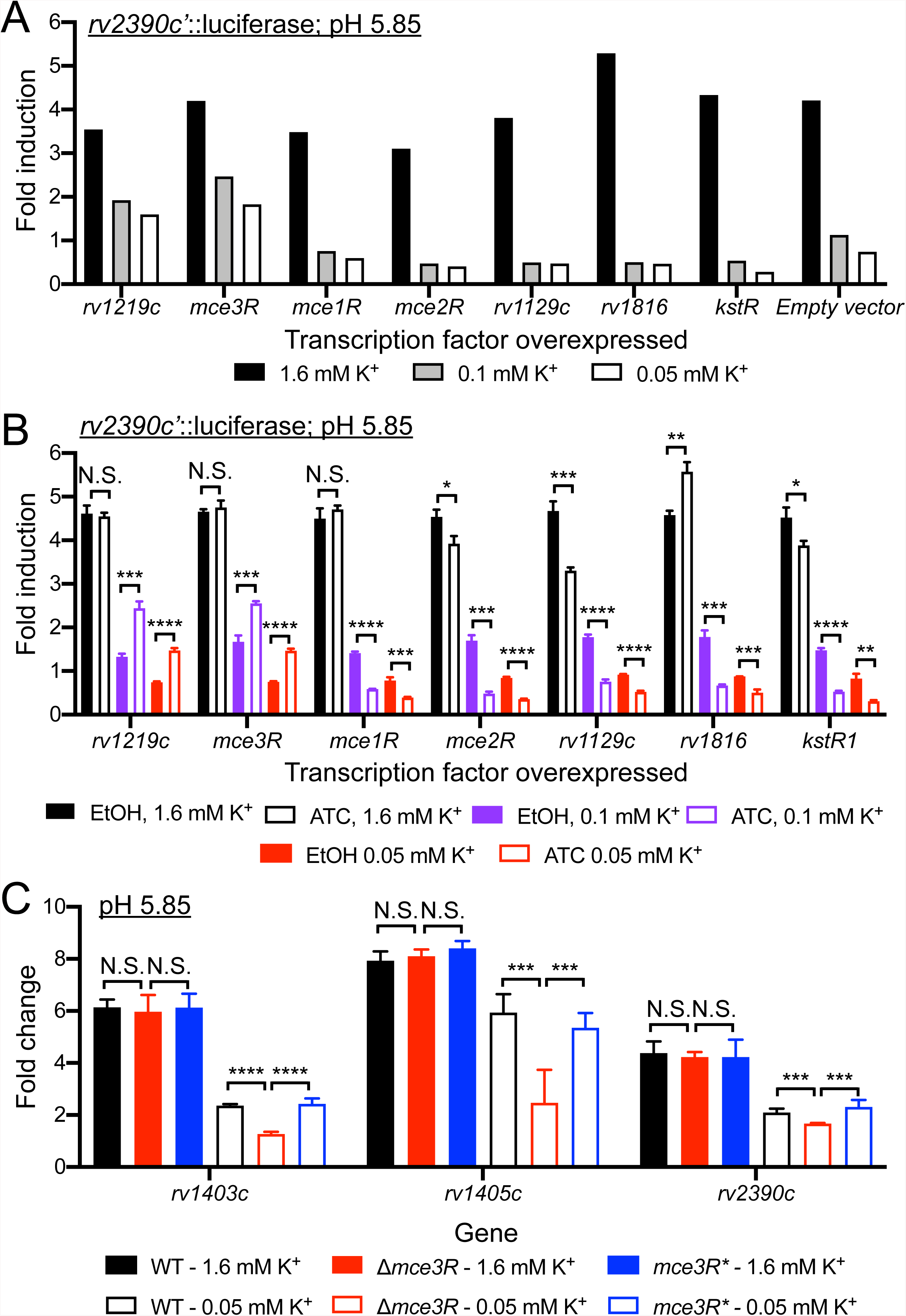
A *rv2390c*’::luciferase reporter transcription factor overexpression screen identifies lipid utilization regulators as modulators of the interplay between environmental [K^+^] and Mtb pH response. (A and B) Lipid utilization regulator hits from reporter-based, inducible transcription factor (TF) overexpression screen. A library of inducible TF overexpression plasmids (P_1_’::*TF*-FLAG-tetON) in the background of a Mtb(*rv2390c*’::luciferase) strain was screened for their response to acidic pH in the presence of low [K^+^]. TF overexpression was induced by adding 200 ng/ml of anhydrotetracycline (ATC) 1 day before Mtb was exposed to K^+^-free 7H9, pH 7 or K^+^-free 7H9, pH 5.85 media supplemented with 1.6 mM or 0.05 mM K^+^. 9 days post-exposure (continuous ATC presence), light output (relative light units, RLU) and OD_600_ were measured. Fold induction compares RLU/OD_600_ in each condition to RLU/OD_600_ in the control K^+^-free 7H9, pH 7 condition. (A) shows results of the lipid utilization regulator hits, together with an empty vector plasmid control. (B) shows validation of the screen hits in (A), with each hit TF compared to its uninduced control (ethanol, “EtOH”, as a carrier control). In (B), data are shown as means ± SD from three experiments, and p-values were obtained with an unpaired t-test with Welch’s correction. N.S. not significant, * p<0.05, ** p<0.01, *** p<0.001, **** p<0.0001. (C) Mce3R modulates the interplay between environmental [K^+^] and Mtb pH response. Log-phase WT, Δ*mce3R* and *mce3R\** (complemented mutant) were exposed for 4 hours to K^+^-free 7H9, pH 7 or K^+^-free 7H9, pH 5.85 media supplemented with 1.6 mM or 0.05 mM K^+^, before RNA was extracted for qRT-PCR analysis. Fold change is as compared to the K^+^-free 7H9, pH 7 condition. *sigA* was used as the control gene, and data are shown as means ± SD from 3 technical replicates, representative of 3 experiments. p-values were obtained with an unpaired t-test with Welch’s correction and Holm-Sidak multiple comparisons, N.S. not significant, *** p<0.001, **** p<0.0001.

We decided to pursue here further study of Mce3R, given its still poorly understood role in Mtb biology. A deletion mutant strain of *mce3R* was generated and tested in the same experimental conditions as in the original *rv2390c’*::luciferase transcription factor overexpression screen. As shown in Figure 2C, in contrast to overexpression of *mce3R* which partially rescued the Mtb pH response phenotype in the presence of low [K⁺], deletion of *mce3R* resulted in the converse phenotype, with a significant reduction in pH regulon gene expression in the mutant strain in acidic pH + K^+^-limiting conditions, compared to the parental wild type (WT) strain. Complementation of the Δ*mce3R* strain restored gene induction to WT levels (Figure 2C).

Taken together, these results indicate that factors related to lipid metabolism regulation play a role in modulating the interplay between external K^+^ levels and the response of Mtb to the critical environmental cue of acidic pH.

### Mtb transcriptional response to acidic pH, low [K^+^], and cholesterol are linked

Given the intriguing connection revealed by the screen between Mtb lipid metabolism regulation, acidic pH response, and environmental [K⁺], we next pursued investigation of how these three facets might be related at the transcriptional level. Examination of the impact of K⁺ limitation on the transcriptional response of Mtb to cholesterol showed that it also dampened induction of the five tested Mtb cholesterol regulon genes (Figure 3A). Interestingly, we also observed a reverse association, where cholesterol further induced the expression of genes in the K⁺ regulon, in the presence of K^+^-limiting conditions (Figure 3B). These data relate the presence of cholesterol to environmental K⁺, further supporting a complex interplay between lipid metabolism, environmental K⁺ levels, and Mtb response to acidic pH.

**Figure 3.**
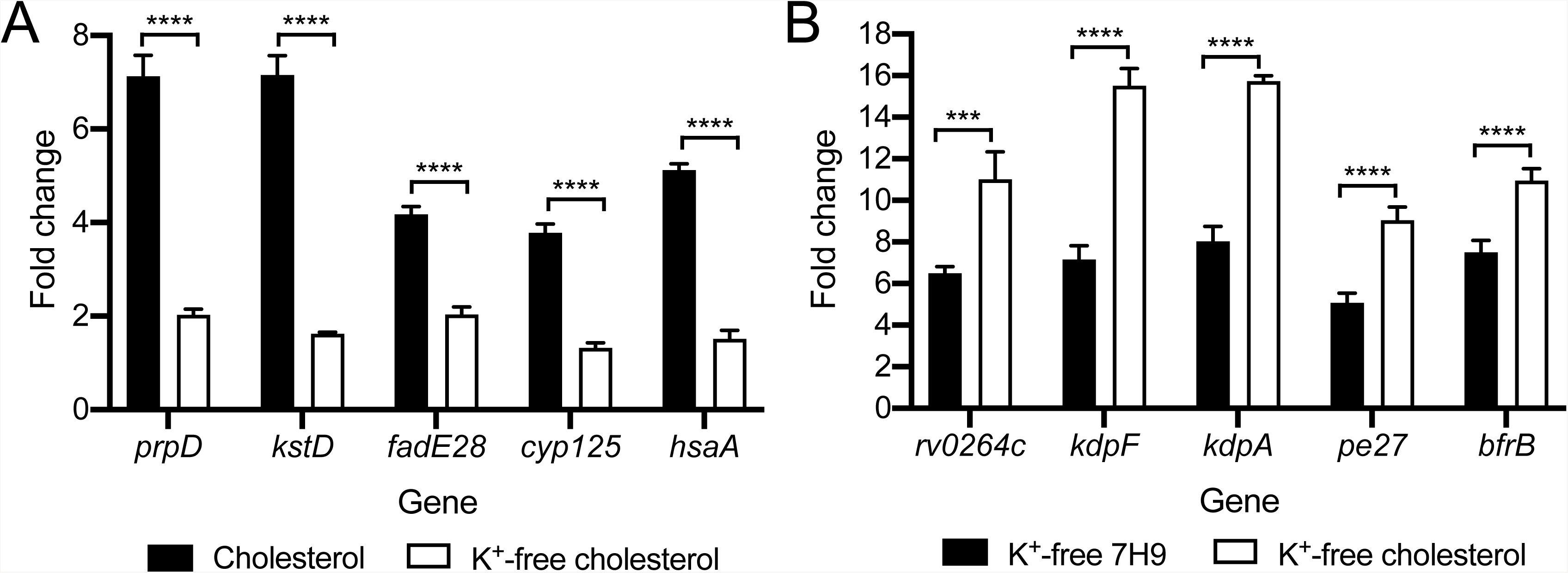
Mtb transcriptional response to low [K^+^] and cholesterol are linked. (A) Low [K^+^] dampens Mtb response to cholesterol. Log-phase Mtb was exposed for 4 hours to (i) 7H9, pH 7, (ii) cholesterol, pH 7, or (iii) K^+^-free cholesterol, pH 7 media, before RNA was extracted for qRT-PCR analysis. (B) Cholesterol augments Mtb response to low [K^+^]. Log-phase Mtb was exposed for 4 hours to (i) 7H9, pH 7, (ii) K^+^-free 7H9, pH 7, or (iii) K^+^-free cholesterol, pH 7 media, before RNA was extracted for qRT-PCR analysis. Fold change is as compared to the 7H9, pH 7 condition in all cases. *sigA* was used as the control gene, and data are shown as means ± SD from 3 technical replicates, representative of 3 experiments. p-values were obtained with an unpaired t-test with Welch’s correction, *** p<0.001, **** p<0.0001.

To obtain a comprehensive understanding of the interplay between the transcriptional response of Mtb to acidic pH and cholesterol, we conducted genome-wide RNA sequencing (RNAseq) analysis, with Mtb exposed to conditions of: (i) 7H9, pH 7 (control), (ii) 7H9, pH 5.7, (iii) cholesterol, pH 7, and (iv) cholesterol, pH 5.7 media. Examination of the impact of acidic pH on the response of Mtb to cholesterol showed an augmentation in the induction of a subset of genes, with expression of 25 differentially expressed genes increased (genes differentially expressed log_2_-fold change ≥1 in the cholesterol, pH 7 condition; log_2_-fold change ≥0.6 between cholesterol, pH 5.7 and cholesterol, pH 7 conditions; p<0.05, FDR<0.01 in both sets) (Figure 4A, Table S2). Further analysis by qRT-PCR indicated a stepwise concentration dependence in this relationship, with expression levels of the cholesterol regulon genes tested showing distinct increase levels with increasing acidity (Figure 4B). On the other hand, increasing the concentration of cholesterol smoothly increased induction levels of the tested genes in both pH 7 and pH 5.7 conditions (Figure 4C).

**Figure 4.**
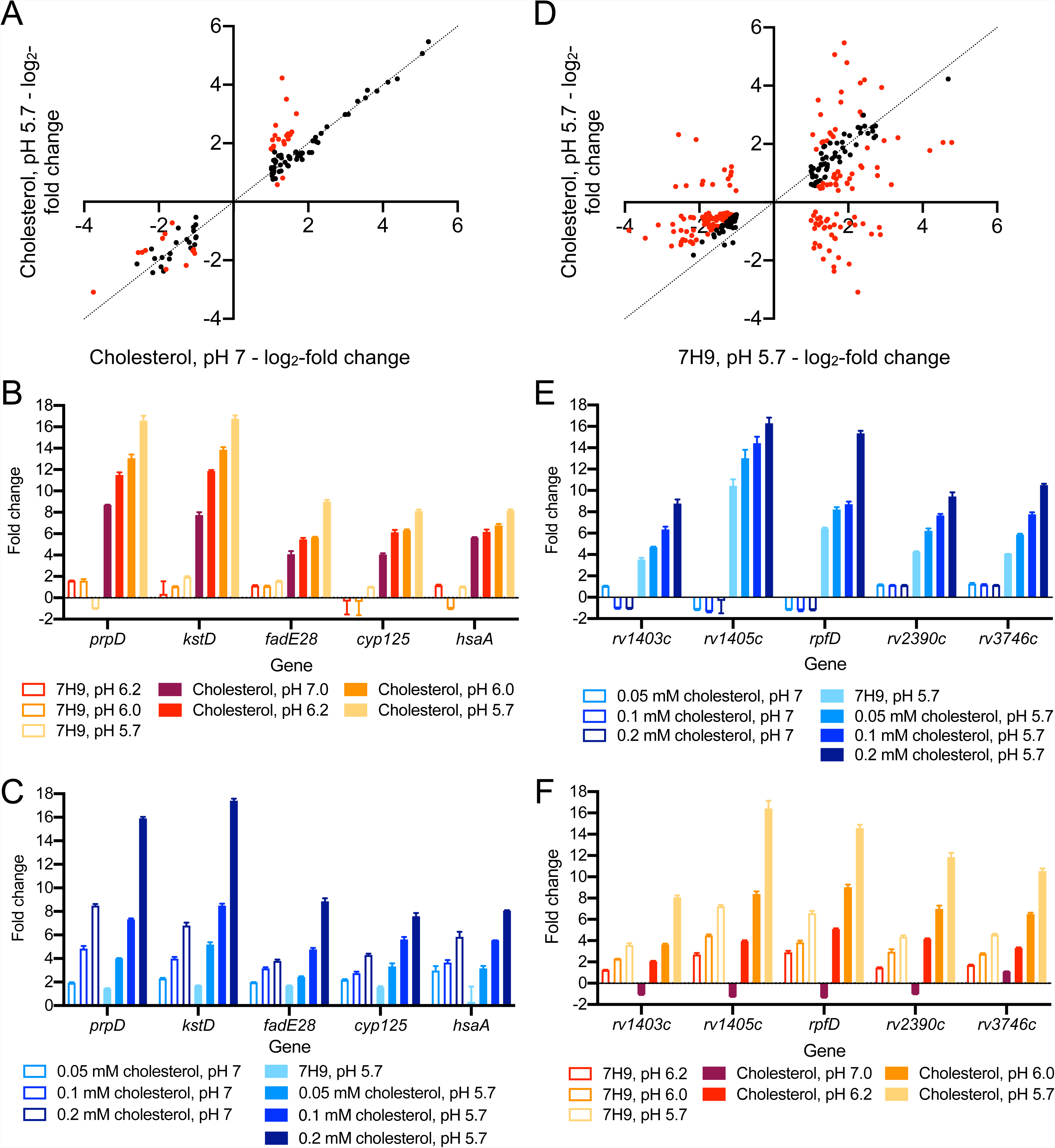
Mtb transcriptional response to acidic pH and cholesterol are linked globally. (A and D) Global changes in Mtb response to cholesterol and acidic pH in the simultaneous presence of both signals. Log-phase Mtb was exposed for 4 hours to (i) 7H9, pH 7, (ii) 7H9, pH 5.7, (iii) cholesterol, pH 7, or (iv) cholesterol, pH 5.7 media, before RNA was extracted for RNAseq analysis. Log_2_-fold change compares gene expression in each indicated condition to the 7H9, pH 7 condition. Genes marked in red had a log2-fold change ≥ 0.6 between the two conditions compared (p<0.05, FDR<0.01 in both sets, with log_2_-fold change ≥1 in the single condition set). (B and C) Effect of acidic pH on Mtb response to cholesterol is concentration dependent. Log-phase Mtb was exposed to the indicated conditions, along with control 7H9, pH7 condition, for 4 hours, before RNA was extracted for qRT-PCR analysis. (E and F) Effect of cholesterol on Mtb response to acidic pH is concentration dependent. For all qRT-PCR data, fold change is as compared to the 7H9, pH 7 condition, and data are shown as means ± SD from 3 technical replicates, representative of 3 experiments.

Complementarily, the RNAseq data showed that the presence of cholesterol also affected differential gene expression of Mtb in response to acidic pH (Figure 4D, Table S3). In this case, the presence of cholesterol increased expression of a subset of pH regulon genes, with expression of 118 differentially expressed gene increased (genes differentially expressed log_2_-fold change ≥1 in the pH 5.7 condition; log_2_-fold change ≥0.6 between cholesterol, pH 5.7 and 7H9, pH 5.7 conditions; p<0.05, FDR<0.01 in both sets), but also interestingly reduced expression of a different subset of genes, with 67 differentially expressed gene decreased (gene differentially expressed log_2_-fold change ≥1 in the pH 5.7 condition; log_2_-fold change ≥0.6 between cholesterol, pH 5.7 and 7H9, pH 5.7 conditions; p<0.05, FDR<0.01 in both sets) (Figure 4D, Table S3). Among this latter group, further examination of the RNAseq data showed that the majority of genes (35/43) most strongly affected (log_2_-fold change ≥1.59 between cholesterol, pH 5.7 and 7H9, pH 5.7 conditions) also exhibited lower expression in the cholesterol, pH 7 condition compared to 7H9, pH 7. As such, the presence of cholesterol alone downregulates expression of these genes. This finding indicates an overlap in a subset of the cholesterol and pH regulon in Mtb; expression of these genes is regulated in opposite directions by the two signals when present individually (induced by acidic pH, repressed by cholesterol), with cholesterol appearing to be the dominant signal when Mtb is exposed to both cues concurrently.

Focusing on genes in the acidic pH regulon that are not also differentially expressed in the presence of cholesterol alone, qRT-PCR analysis showed a concentration dependence in the augmentation of gene expression induction at acidic pH with increasing cholesterol concentration (Figure 4E). pH level-dependence of induction of the acidic pH regulon genes tested was also observed in both standard 7H9 and cholesterol conditions (Figure 4F). Overall, these findings show the global cross-regulation between cholesterol and pH response in Mtb, with the concurrent presence of each factor strongly influencing the expression of genes in each regulon.

### Mce3R regulates Mtb response to cholesterol only in the context of acidic pH

Our results above raise the question of how Mtb response to acidic pH and cholesterol is cross-regulated. To gain insight into possible regulatory mechanisms underlying the synergy in Mtb transcriptional response to cholesterol and acidic pH in the concurrent presence of both cues, we first tested the effect of a *phoPR* deletion on these responses. PhoPR is a well-established two-component system known to play a crucial role in Mtb response to the linked environmental cues of acidic pH and Cl^−^^1,2^. Consistent with this, we observed that a Δ*phoPR* mutant exhibited significantly reduced induction of acidic pH regulon genes in both standard 7H9 or cholesterol media as a base (Figure S1A). Notably however, the simultaneous presence of cholesterol still resulted in an increase in the induction level of several tested genes, compared to the single acidic pH condition, in Δ*phoPR* Mtb (Figure S1A, compare open red bars to open black bars). The same phenomenon of an overall decrease in gene expression induction with the Δ*phoPR* Mtb mutant in both standard 7H9 or cholesterol media as a base was also observed with Cl^−^ as the inducing signal instead of acidic pH (Figure S1B). Similarly, the continued increase in gene expression induction for a few tested genes in the simultaneous presence of cholesterol was also observed in the dual cholesterol/high [Cl^−^] condition compared to the single high [Cl^−^] condition (Figure S1B, compare open blue bars to open black bars). Complementation of the Δ*phoPR* Mtb mutant restored gene expression profiles to WT levels in all cases (Figure S1). These findings suggest the involvement of additional regulators that contribute to the coordination of the acidic pH/Cl^−^ response with Mtb response to cholesterol.

The putative lipid metabolism regulator Mce3R had been found as a robust hit in the transcription factor overexpression library screen described above, which first revealed the ability of lipid metabolism regulators to modulate the interplay between environmental K^+^ levels and the response of Mtb to acidic pH. While Mce3R is known to regulate the *mce3* operon and associated upstream genes^31–33^, its possible role in the overall response of Mtb to cholesterol is unknown. To determine if Mce3R might also play a role in coordinating Mtb acidic pH and cholesterol response, we performed global RNAseq analysis on the Δ*mce3R* mutant and compared the transcriptional profiles obtained to those of WT Mtb. The same four conditions utilized above for WT Mtb were used here, namely: (i) 7H9, pH 7 (control), (ii) 7H9, pH 5.7, (iii) cholesterol, pH 7, and (iv) cholesterol, pH 5.7 media. A first observation was that *mce3R* deletion had little effect on global transcriptional response of Mtb to acidic pH or cholesterol when either condition was present alone (Figures 5A and 5B, Tables S4 and S5). Strikingly however, in the dual cholesterol, pH 5.7 condition, a subset of genes associated with the cholesterol regulon were significantly less induced in the Δ*mce3R* Mtb mutant compared to WT (Figure 5C, Table S6). Follow-up qRT-PCR experiments validated that *mce3R* deletion only affected expression of cholesterol regulon genes in Mtb in the context of the simultaneous presence of cholesterol and acidic pH, with complementation restoring gene expression to WT levels (Figures 5D and 5E). Markedly, for the Δ*mce3R* mutant, the induction levels of the cholesterol regulon genes in the dual cholesterol, pH 5.7 condition were very similarly to those observed in the single cholesterol (pH 7) condition (compare open bars in Figure 5E to open bars in Figure 5D). Of note, augmentation in expression of acidic pH regulon genes by cholesterol was however still observed in the Δ*mce3R* mutant as compared to WT Mtb (Figure 5F).

**Figure 5.**
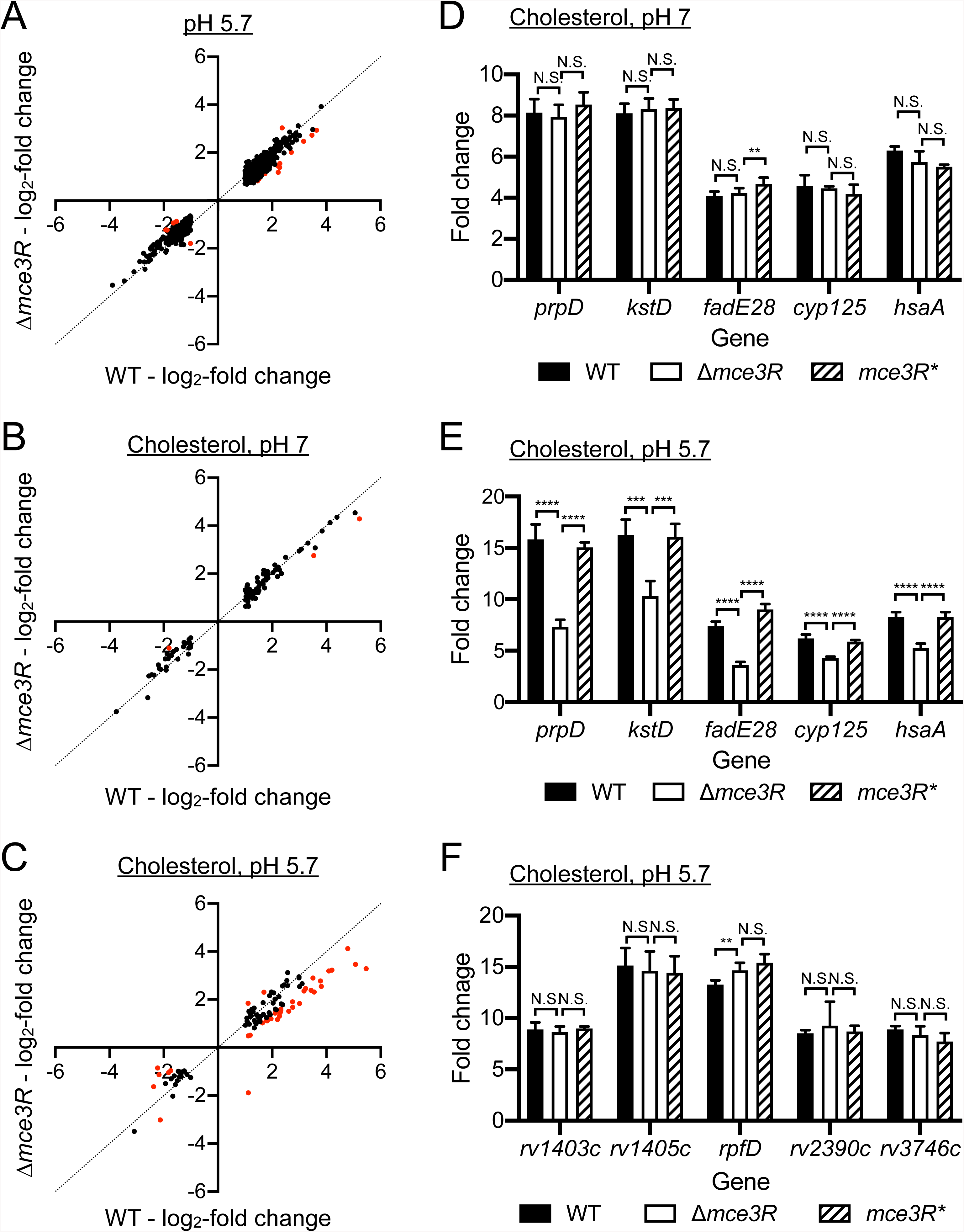
Mce3R regulates Mtb response to cholesterol only in the context of acidic pH. Log-phase WT, Δ*mce3R* and *mce3R*\* (complemented strain) Mtb were exposed for 4 hours to (i) 7H9, pH 7, (ii) 7H9, pH 5.7, (iii) cholesterol, pH 7, or (iv) cholesterol, pH 5.7 media, before RNA was extracted for RNAseq (A-C) or qRT-PCR (D-F). For RNAseq data, log_2_-fold change compares gene expression in each indicated condition to the 7H9, pH 7 condition. Genes marked in red had a log2-fold change ≥ 0.6 between Δ*mce3R* and WT strains in the indicated condition (p<0.05, FDR<0.01 in both sets, with log_2_-fold change ≥1 in WT). For qRT-PCR data, fold change is as compared to the 7H9, pH 7 condition. Data are shown as means ± SD from 3 technical replicates, representative of 3 independent experiments. p-values were obtained with as unpaired t-test with Welch’s correction and Holm-Sidak multiple comparisons. N.S. not significant, ** p<0.01, *** p<0.001, **** p<0.0001.

Given (i) the results above, (ii) our finding that cholesterol also augments Mtb response to low [K^+^] (Figure 3B), and (iii) the original identification from our screen of Mce3R as a regulator able to modulate the interplay between environmental K^+^ levels and the response of Mtb to acidic pH, we next sought to test if Mce3R would also have a role in coordinating Mtb response in the simultaneous presence of both cholesterol and low [K^+^]. WT, Δ*mce3R*, and the complemented Δ*mce3R* Mtb mutant (*mce3R\**) were exposed to: (i) K⁺-free 7H9, pH 7, or (ii) K⁺-free cholesterol, pH 7 media, for 4 hours before RNA were extracted for gene expression analysis. Strikingly, the ability of cholesterol to augment induction of the K^+^ regulon genes was largely lost in the Δ*mce3R* mutant, even as deletion of *mce3R* had no effect on induction of the K^+^ regulon genes in the absence of cholesterol (Figure 6A). The dampening of cholesterol gene induction by the simultaneous presence of low [K^+^] was additionally unchanged by deletion of *mce3R* (Figure 6B).

**Figure 6.**
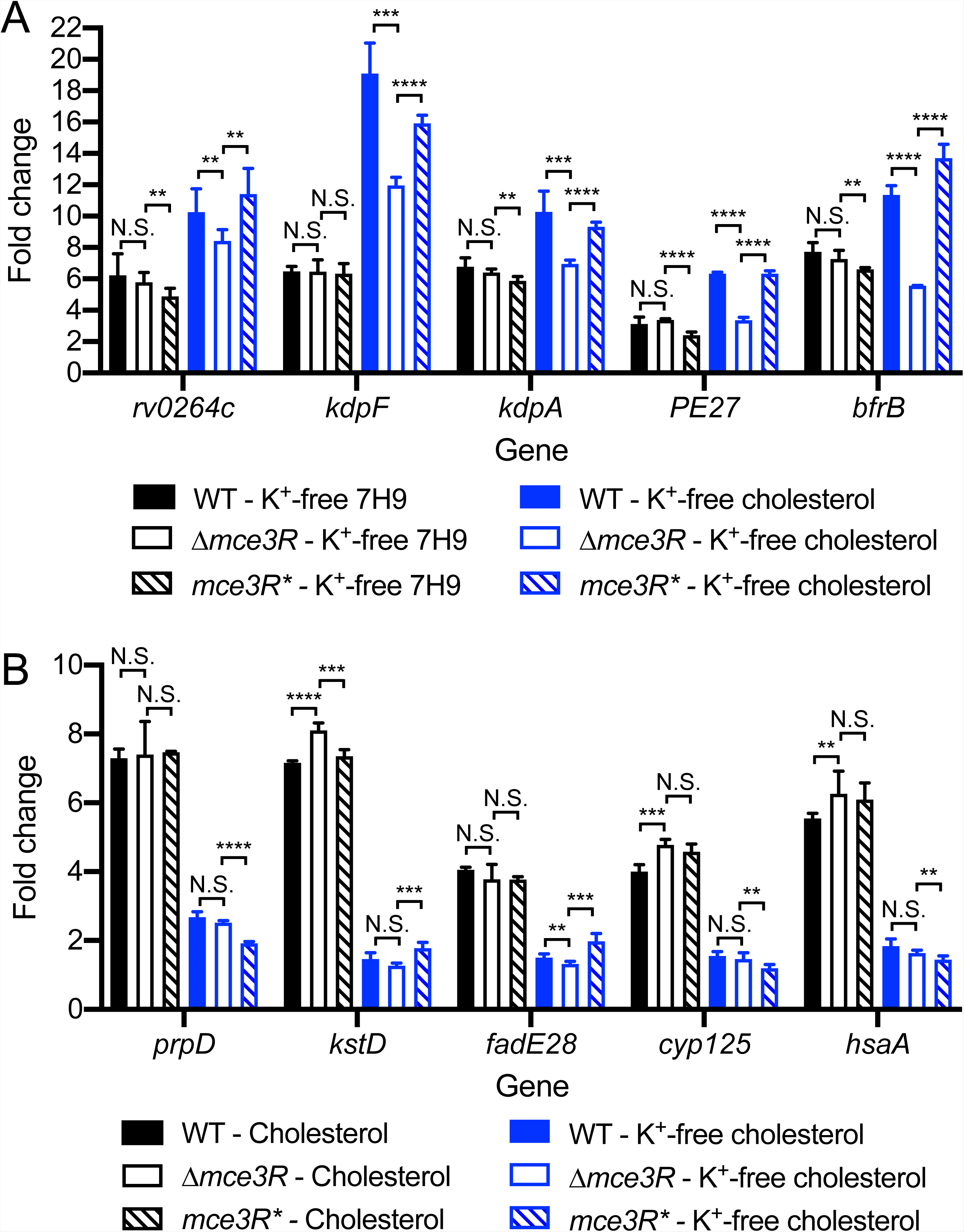
Mce3R deletion dampens Mtb response to K^+^ in the presence of cholesterol. Log-phase WT, Δ*mce3R*, and *mce3R*\* (complemented strain) Mtb were exposed for 4 hours to (A) K^+^-free 7H9 or cholesterol media at pH 7, or (B) cholesterol or K^+^-free cholesterol media at pH 7, along with 7H9, pH 7 as the control condition. Fold change is as compared to the 7H9, pH 7 condition in all cases. *sigA* was used as the control gene, and data are shown as means ± SD from 3 technical replicates, representative of 3 experiments. p-values were obtained with an unpaired t-test with Welch’s correction and Holm-Sidak multiple comparisons, N.S. not significant, ** p<0.01, *** p<0.001, **** p<0.0001.

Together, these results excitingly identify Mce3R as a transcription factor that specifically acts to coordinate Mtb response in the presence of at least two environmental signals (cholesterol + acidic pH, low [K^+^] + cholesterol, acidic pH + low [K^+^]), with little effect when only one signal is present. There is further directionality in its activity, as in each case, *mce3R* deletion affected response to only one of the two signals concurrently present. These findings emphasize the interconnectedness between Mtb response to acidic pH, low [K^+^], and cholesterol, and the multifaceted role of Mce3R in the coordinated regulation of Mtb response to its local environment.

### A Δ*mce3R* Mtb mutant is attenuated for host colonization in the presence of lipids

Changes in pH and [K^+^], and differing presence of cholesterol, are all factors expected to occur during Mtb host colonization. To assess the importance of Mce3R during infection, we infected bone marrow-derived macrophages (BMDMs) with WT, Δ*mce3R*, or *mce3R\** Mtb. Given our findings of the importance of Mce3R in coordination of the cholesterol response of Mtb, these infections were also performed in BMDMs pre-treated with oleate to induce the formation of foamy macrophages^35,36^, to determine whether infection in lipid-rich macrophages would differ from untreated BMDMs. As shown in Figure 7A, the Δ*mce3R* mutant was attenuated for growth in BMDMs compared to the WT and *mce3R\** Mtb, with a more pronounced effect in the context of foamy macrophage infection. While Mtb actively prevents complete maturation of the phagosome, it is still exposed to slightly decreased pH levels during macrophage infection^37^. These results thus support a role of Mce3R in the ability of Mtb to respond appropriately to its local environment for continued growth.

**Figure 7.**
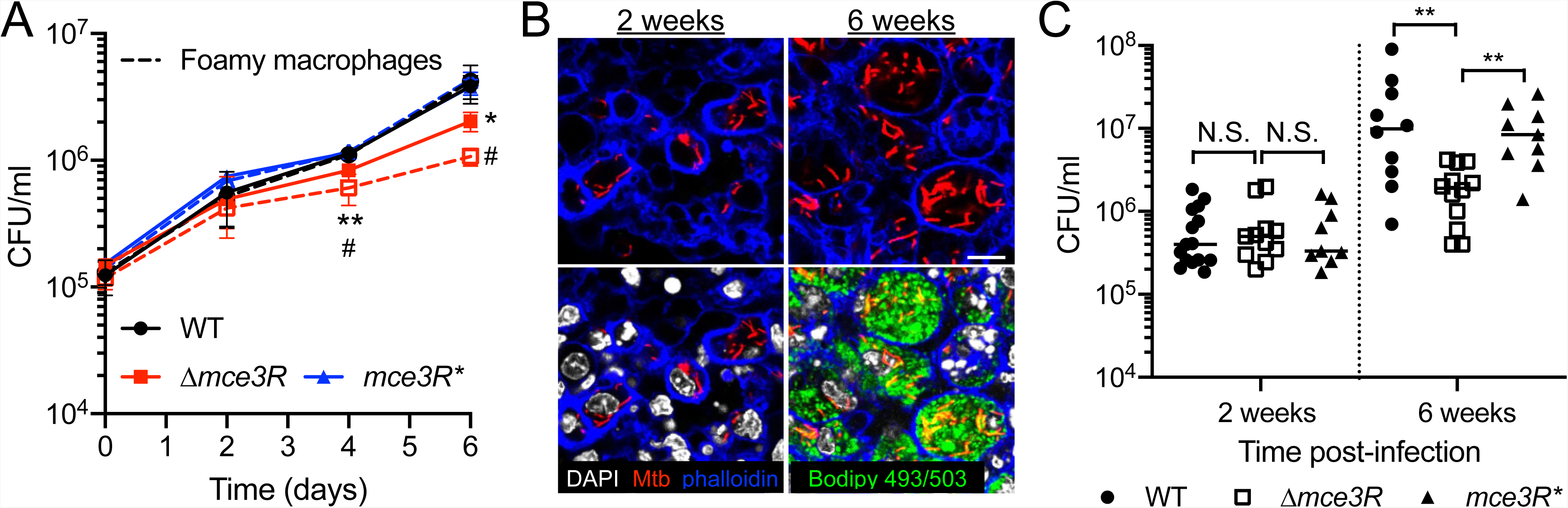
A Δ*mce3R* Mtb mutant is attenuated for host colonization in the presence of lipids. (A) Δ*mce3R* Mtb is attenuated for macrophage colonization. Murine bone marrow-derived macrophages untreated or pre-treated with oleate for 24 hours to induce foamy macrophages were infected with WT, Δ*mce3R*, or *mce3R*\* (complemented mutant) Mtb, and colony forming units (CFUs) tracked over time. Data are shown as means ± SD from 3 wells, representative of 3 independent experiments. p-values were obtained with an unpaired t-test with Welch’s correction, comparing Δ*mce3R* to WT Mtb in untreated macrophages (*) or foamy macrophages (#). */# p<0.05, ** p<0.01. (B) Mtb is present in foamy macrophages 6 weeks post-infection in C3HeB/FeJ mice. C3HeB/FeJ mice were infected with Mtb constitutively expressing mCherry, and animals sacrificed at 2 or 6 weeks post-infection. Lungs were fixed and processed for confocal microscopy imaging. Images shown are z-slices from reconstructed 3D images. Nuclei are shown in grayscale (DAPI), all bacteria are marked in red (mCherry), f-actin is shown in blue (phalloidin), and lipid droplets are shown in green (Bodipy 493/503). Scale bar 10 µm. (C) Δ*mce3R* Mtb is attenuated for colonization in a murine infection model. C3HeB/FeJ mice were infected with WT, Δ*mce3R*, or *mce3R*\* Mtb, and lung homogenates plated for CFUs 2 or 6 weeks post-infection. p-values were obtained with a Mann-Whitney statistical test. N.S. not significant, **p<0.01.

Finally, we examined the role of Mce3R in Mtb infection of a whole animal host using the C3HeB/FeJ murine model. Importantly, this mouse strain recapitulates key lesion types observed during human infection^38–40^. Of pertinence here, foamy macrophages are present at 6 weeks post-infection, both in macrophage-rich lesions and in the cuff surrounding caseous necrotic lesions that are the hallmark of tuberculosis disease (Figure 7B)^41^. The mice were infected with WT, Δ*mce3R*, or *mce3R\** Mtb, and the infection allowed to progress for 2 or 6 weeks. We did not observe significant differences in bacterial load among the three strains at the 2-week time point (Figure 7C), consistent with this time point being prior to the formation of foamy macrophages (Figure 7B). In contrast, at the 6-week time point, the Δ*mce3R* mutant exhibited a significantly reduced bacterial load compared to the WT and *mce3R\** Mtb strains (Figure 7C).

Together, these results demonstrate that Mce3R plays an important role in Mtb growth in macrophages, with the absence of Mce3R resulting in a significantly reduced ability of Mtb to maintain persistent colonization of a whole animal host.

## DISCUSSION

From the alveolar airspace to the macrophage phagosome to the granuloma, Mtb encounters a complex and dynamic environmental milieu that plays a crucial role in shaping its infection biology. This milieu encompasses signals originating from the cellular and tissue microenvironment, which often reflect the host’s immune response, and changes to available nutrient sources. There have been extensive studies seeking to understand the response of Mtb to specific environmental signals and nutrients, and the regulation underlying such responses. In contrast, significantly less is known regarding the interplay between Mtb response to particular environmental stimuli and available nutrients, and the regulatory mechanisms that enable the integration of such responses. Our study here reveals the intrinsic relationships between Mtb response to low [K^+^], acidic pH, and cholesterol, with synergy in the acidic pH and cholesterol response, and dampening of Mtb response to both signals in the presence of low [K^+^]. Concurrent exposure to acidic pH and cholesterol is expected to be a critical aspect of the local environment during Mtb infection, in the context of bacterial residence in phagosomes of foamy macrophages, a major host cell type observed not just in animal models of infection, but also in human infections^38,39,42,43^. Indeed, Mtb growth in acidic pH media is improved when cholesterol is provided as the carbon source^18^, and our finding of synergy in Mtb transcriptional response to these two signals provides a crucial new dimension to the understanding of Mtb adaptive biology in this environment.

Changes in expression of bacterial “virulence factors”, driven by changes in the presence of specific metabolites, have been increasingly described for different bacterial pathogens^44^. This includes increased expression of leucocidins in *Staphylococcus aureus* in the presence of pyruvate^45^, and induced expression of *Salmonella* pathogenicity island 2 genes in the presence of succinate^46^. Our results demonstrating the reciprocal augmentation of Mtb response to acidic pH and cholesterol in the concurrent presence of both signals, and the increased expression of K^+^-responsive genes in the simultaneous presence of cholesterol, expands on this facet of bacterial biology. In particular, together with previous studies that have shown increased transcription of genes in the Cl^−^ regulon in the simultaneous presence of cholesterol^20^, and increased expression of cholesterol utilization genes in iron-limiting conditions^21,22^, our findings support the existence of an intrinsic interplay between Mtb response to environmental cues and the critical nutrient of cholesterol.

Strikingly, we identify Mce3R as a transcription factor that critically functions in the integration of these signals. Specifically, a Δ*mce3R* mutant fails to synergistically upregulate genes in the cholesterol regulon in the simultaneous presence of acidic pH, and in the K^+^ regulon in the simultaneous presence of cholesterol, even while response to each signal individually is unaffected. It will be intriguing in future studies to explore the underlying mechanism enabling the triggering of Mce3R function only in the context of the concurrent presence of two signals versus one. Mce3R is a cytoplasmic protein and would not directly sense changes in external signals, although an increase in cholesterol utilization would result in increased levels of cholesterol within the bacterium. Of note, the affinity of Mce2R and Mce1R for binding to their target DNA have been reported (experimentally or computationally respectively) to change in the presence of long-chain fatty acids^25,47^. Whether and how Mce3R binding to DNA may be affected by cholesterol, and whether interacting partners exist that aid in transducing the external stimuli to Mce3R, are thus areas for future investigation. In addition, a recent report indicated that a Δ*mce3R* Mtb mutant was less susceptible to oxidative stress^48^, a host defense response also encountered by the bacteria during infection of phagocytes^49^. Further studies are required to determine if Mce3R may also act in integrating oxidative stress signals with cholesterol response in Mtb.

As in mammalian cells, K^+^ is also the most abundant cation in bacterial cells, and the importance of K^+^ in host colonization is becoming increasingly appreciated^3,50,51^. Our unexpected finding linking Mtb lipid utilization regulators to the bacterium’s K^+^ response opens a new facet of study in understanding how K^+^ response/homeostasis integrates with the metabolism of a bacterium. Intriguingly, cholesterol regulation of K^+^ channel activity has been demonstrated in mammalian systems, with the inwardly rectifying K^+^ (Kir) channels shown to possess cholesterol-binding sites^52,53^. Regulation of the prokaryotic homolog of Kir (KirBac1.1) by cholesterol has also been demonstrated^54^. Cholesterol binding usually downregulates Kir channel function, although it has been shown to enhance activity of certain Kir channels, such as Kir3.4^55^. In Mtb, the CeoBC K^+^ uptake system is critical in maintaining K^+^ homeostasis^3,56^. Whether cholesterol directly regulates CeoBC activity in Mtb, and more broadly K^+^ uptake systems in other pathogens, is an interesting question for future study.

Exposure to different environmental signals with a simultaneous change in available nutritional source is a widespread phenomenon for bacterial pathogens and not unique to Mtb. We propose that further study of bacterial responses to concurrent environmental and nutritional cues will yield important new insight into bacterial signal integration and their underlying mechanisms. Unique nodes represented by regulatory factors that specifically act to coordinate bacterial environmental response to multiple signals and thus adaptation, such as Mce3R as identified here, further represent exciting new candidates for therapeutic targeting^57^.

## ACKNOWLEDGEMENTS

We thank members of the Tan laboratory for helpful discussion, and Trever Smith for initial assistance with RNA sequencing analysis. This work was supported by grants from the National Institutes of Health (R01 AI143768 and R21 AI171356) to ST, and by a Natalie V. Zucker Research Center for Women Scholars Research Award from Tufts University School of Medicine to YC.

## AUTHOR CONTRIBUTIONS

Conceptualization, YC. and ST.; Investigation, YC, NJM, and ST; Writing – original draft, YC and ST; Writing – review and editing, YC, NJM., and ST; Supervision, ST.; Funding acquisition, YC and ST.

## STAR ★ Methods

### Key resources table

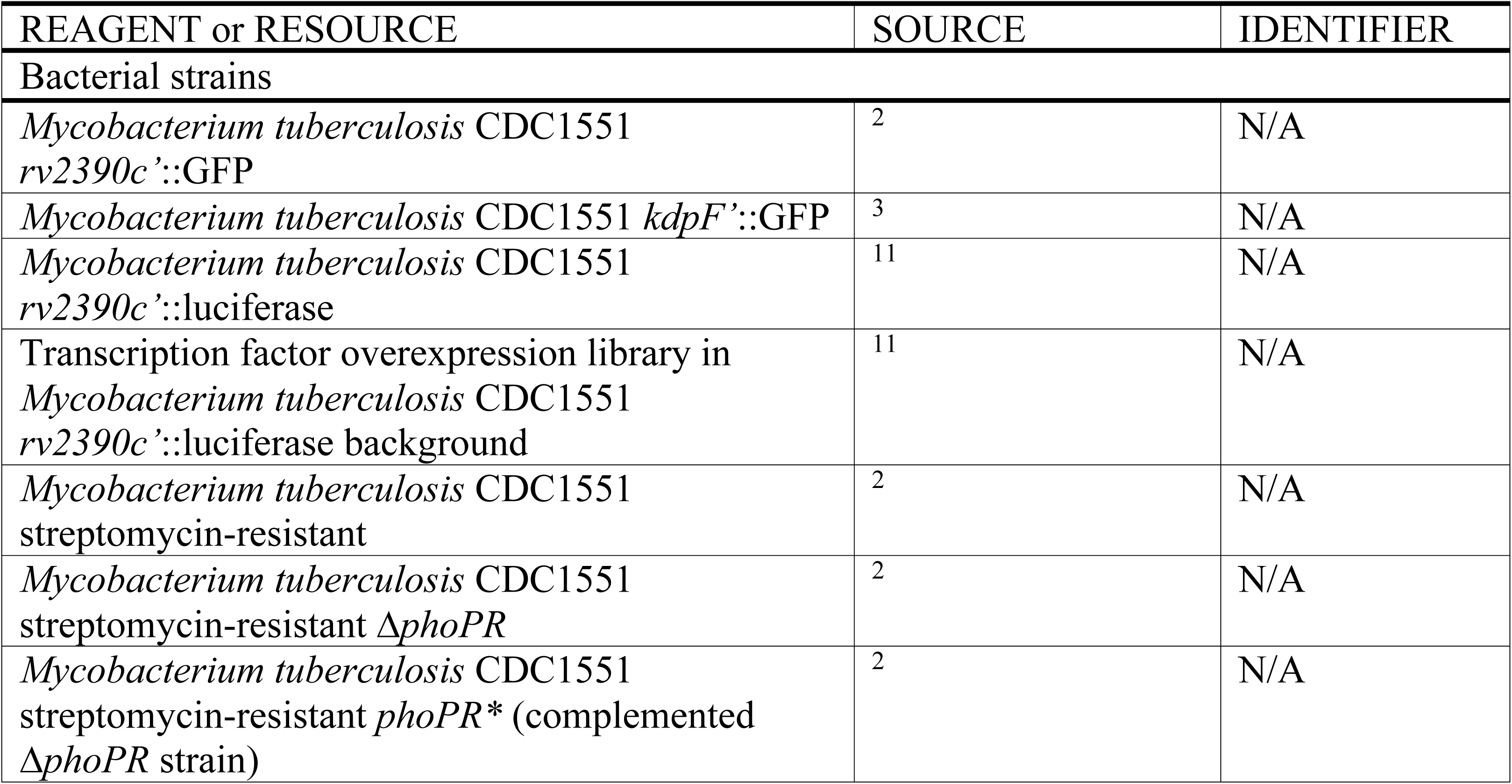
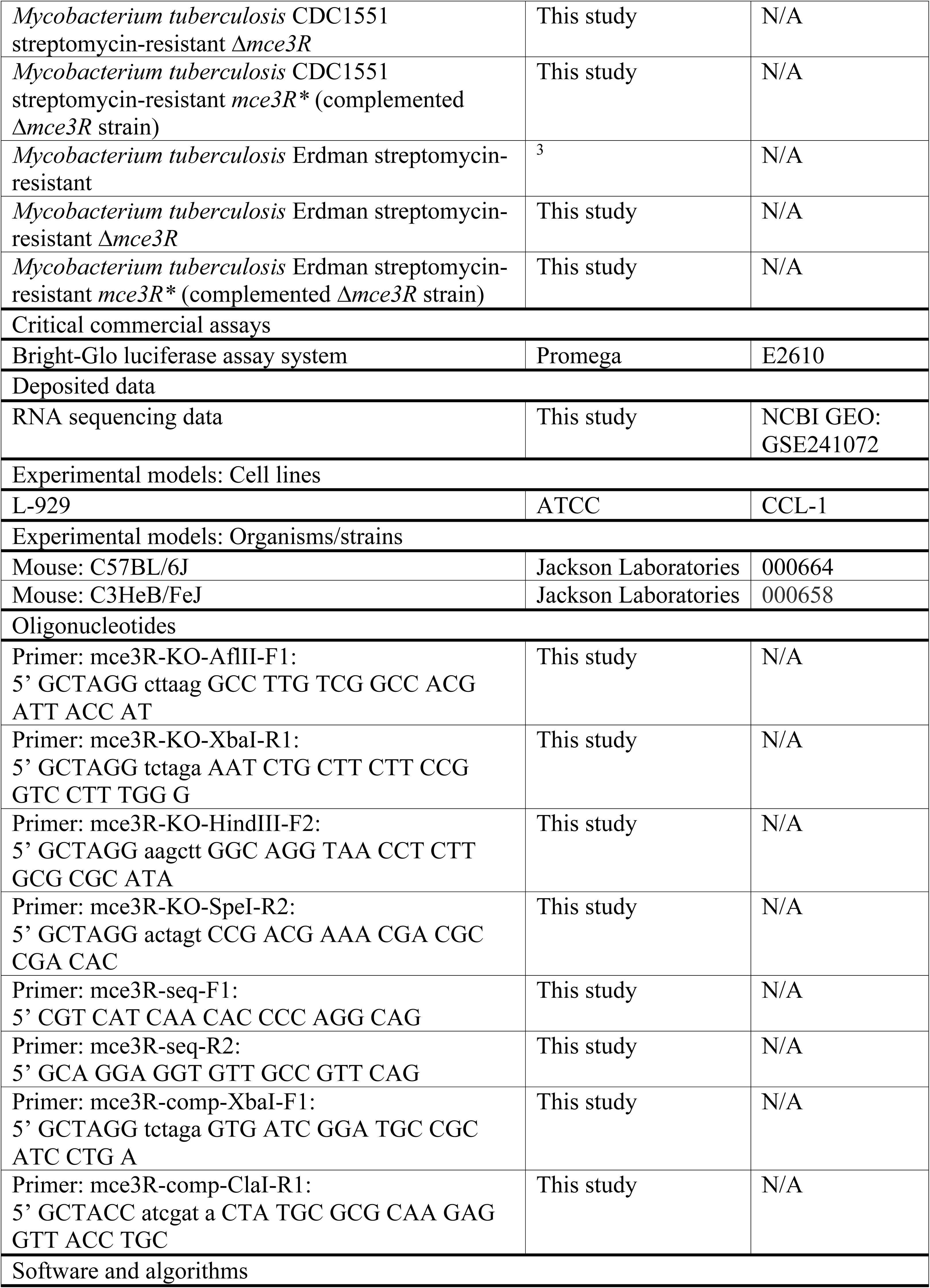
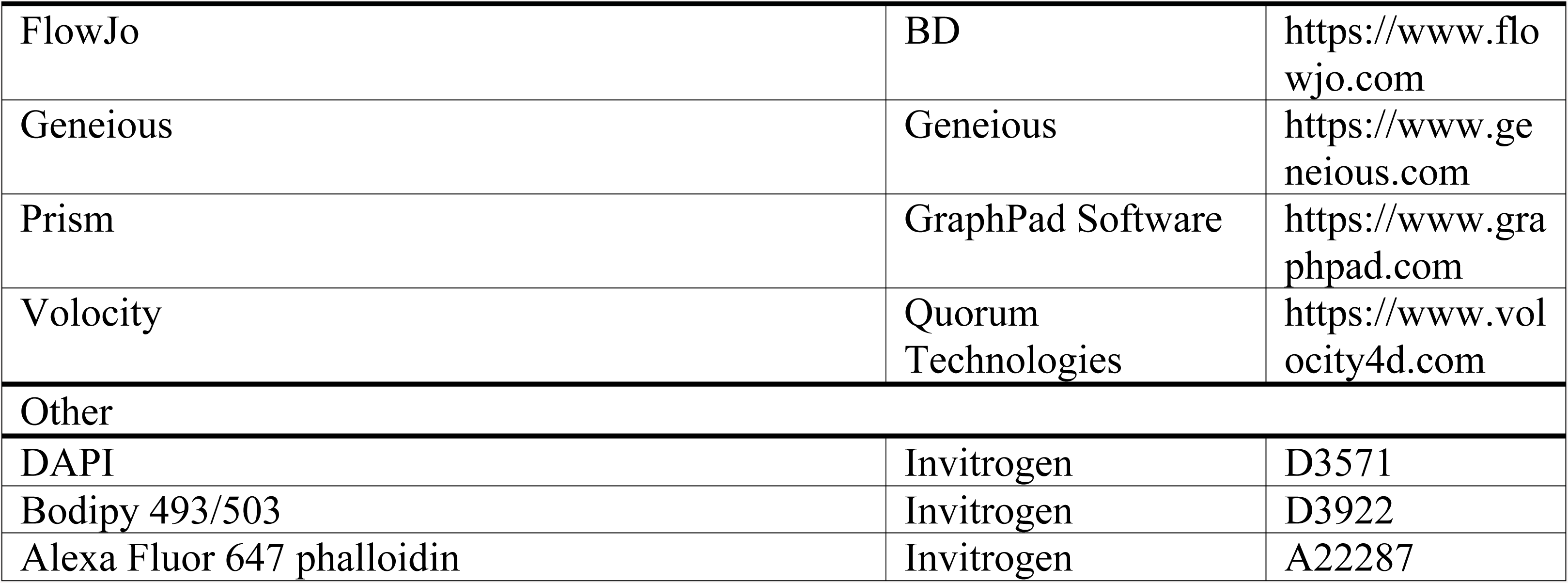

### Resource availability

Further information and requests for resources and reagents should be directed to and will be fulfilled by the lead contact, Shumin Tan.

### Materials availability

All newly generated Mtb strains are available on request from the lead contact, to investigators with the necessary biosafety level 3 facilities to receive and work with these materials.

### Data and code availability

All RNA sequencing data have been deposited in the NCBI GEO database (GSE241072).
This paper does not report original code.
Any additional information required to reanalyze the data reported in this work paper is available from the lead contact upon request

### Experimental model and subject details

#### Murine strains

C57BL/6J and C3HeB/FeJ wild type mice were obtained from Jackson Laboratories. All mice used were female and aged 4 weeks on arrival. Mtb infections were carried out when the mice were 6 weeks of age. All animal protocols in this research followed the guidelines from The National Institutes of Health “Guide for Care and Use of Laboratory Animals”. All animal protocols (#B2021-139) were reviewed and approved by the Institutional Animal Care and Use Committee at Tufts University, in accordance with guidelines from the Association for Assessment and Accreditation of Laboratory Animal Care, the US Department of Agriculture, and the US Public Health Service.

### Method details

#### Mtb strains and culture

Mtb cultures were propagated and maintained as previously described^58^. K^+^-free and cholesterol media were prepared as previously described^3,20^. All antibiotics were added as appropriate at the following concentrations: 100 µg/ml streptomycin, 50 µg/ml hygromycin, 50 µg/ml apramycin, and 25 µg/ml kanamycin. Strains for *in vitro* assays were in the CDC1551 background, while infection assays were carried out with strains in the Erdman background. The *rv2390c’*::GFP, *kdpF’*::GFP, and *smyc’*::mCherry reporters, and the Δ*phoPR* strain and its complement have all been previously described^2,3^. The Δ*mce3R* mutant and its complement were constructed as previously described^2^, with the Δ*mce3R* mutation consisting of a deletion beginning at the *mce3R* start codon (as annotated in the Erdman Mtb strain) through 26 bp from the *mce3R* stop codon. The *rv2390c’*::luciferase transcription factor overexpression library has been previously described^11^.

#### Reporter Mtb strain broth assays

Reporter strains at log-phase were subcultured to an OD_600_ = 0.05 and resuspended in the appropriate media. These were: (i) 7H9, pH 7, (ii) 7H9, pH 5.85, and (iii) K^+^-free 7H9 supplemented with 0.1 mM K^+^, pH 5.85, for assays with the *rv2390c’*::GFP reporter. For assays with the *kdpF’*::GFP reporter, media conditions were: (i) 7H9, pH 7, (ii) 7H9, pH 5.7, (iii) K^+^-free 7H9, pH 7, and (iv) K^+^-free 7H9, pH 5.7. For broth assays conducted in K^+^-free media, an additional wash step with K^+^-free 7H9, pH 7 medium was performed prior to resuspending the cultures in the final assay media. At each time point, aliquots of the cultures were taken and fixed in 4% paraformaldehyde (PFA) in phosphate-buffered saline (PBS). Reporter GFP signal was measured using a BD FACSCalibur flow cytometer, and the data analyzed using FlowJo software (BD).

#### *rv2390c’*::luciferase transcription factor overexpression library screen

To screen the *rv2390c’*::luciferase, tetracycline-inducible transcription factor overexpression (TFOE) library, thawed TFOE Mtb strains in 96-well plates (35 µl volume) were mixed with 165 µl of 7H9, pH 7 medium supplemented with 25 µg/ml kanamycin and 50 µg/ml hygromycin. The plates were then incubated for 11 days at 37°C in a 5% CO_2_ incubator. Subsequently, the TFOE *rv2390c’*::luciferase strains were subcultured at a 1:10 dilution into 100 µl of fresh 7H9, pH 7 medium with appropriate antibiotics and incubated for an additional eight days. After eight days of growth, 200 ng/ml anhydrotetracycline (ATC) was added to induce overexpression of each transcription factor for 24 hours. Following the induction, all samples were transferred to a 96-well v-bottom plate (Corning Costar) and pelleted at 3,000 rpm for 10 minutes. The samples were then washed once with K^+^-free 7H9, pH 7. Each strain was then inoculated at a 1:10 dilution into 100 µl of the following media conditions: (i) K^+^-free 7H9, pH 7, supplemented with 0.05 mM KCl, (ii) K^+^-free 7H9, pH 5.85, supplemented with 1.6 mM KCl, (iii) K^+^-free 7H9, pH 5.85, supplemented with 0.1 mM KCl, and (iv) K^+^-free 7H9, pH 5.85, supplemented with 0.05 mM KCl, in clear-bottom white 96-well plates (Corning Costar). The cultures were then incubated for nine days at 37°C in a 5% CO_2_ incubator. All media contained 50 µg/ml hygromycin and 25 µg/ml kanamycin, as well as 200 ng/ml ATC for continued transcription factor overexpression. Following this nine day incubation, luminescence was assessed using the Bright-Glo luciferase assay system (Promega) as previously described, with both the light output (relative light units, RLU), and the OD_600_ measured using a Biotek Synergy Neo2 multi-mode microplate reader^11^. Fold induction was determined by comparing the RLU/OD_600_ values of the samples from each environmental condition to the RLU/OD_600_ value of Mtb in the K^+^-free 7H9, pH 7, 0.05 mM KCl control condition.

#### RNA sequencing and qRT-PCR analyses

For RNA sequencing (RNAseq) and qRT-PCR analyses, log-phase Mtb cultures (OD_600_ ∼0.6) were used to inoculate standing T25 flasks with filter caps at an OD_600_ = 0.3, containing 10 ml of the following media: (i) 7H9, pH 7.0, (ii) 7H9, pH 5.7, (iii) cholesterol, pH 7.0, or (iv) cholesterol, pH 5.7. For transcriptional analysis of Mtb under K^+^-free conditions, an additional wash step with K^+^-free 7H9, pH 7, was performed prior to inoculation of the culture into the various test conditions: (i) K^+^-free 7H9, pH 7, supplemented with 0.05 mM KCl (control), (ii) K^+^-free 7H9, pH 5.85, supplemented with 1.6 mM KCl, (iii) K^+^-free 7H9, pH 5.85, supplemented with 0.1 mM KCl, or (iv) K^+^-free 7H9, pH 5.85, supplemented with 0.05 mM KCl. After four hours of exposure to the specific environmental conditions, Mtb samples were collected and RNA isolated as previously described^1^. For RNAseq, two biological replicates were prepared for each condition. Library preparation was performed by the Tufts University Genomics Core Facility using the Illumina stranded total RNA with Ribo-Zero Plus kit, and barcoded samples were pooled and sequenced on an Illumina HiSeq 2500 (single-end 100 bp reads). RNAseq data were analyzed using the Geneious program (version 2023.0.4). Specifically, raw data were trimmed using BBDuk with the following parameters: trim adapters (All Truseq), Kmer lengh 27, trim low quality on both ends with minimum quality 30, discard short reads with minimum length 36 bp, minimum entropy 0.1, entropy window size 4 and entropy Kmer size 5. Trimmed sequences were assembled onto the CDC1551 genome (GenBank AE000516.2) using the Bowtie2 mapper^59^ with end-to-end alignment. Gene expression levels were then calculated and excluded ambiguously mapped reads from calculation, with comparison of expression levels performed via DEseq2^60^. qRT-PCR experiments were carried out and data analyzed as previously described^3^.

#### Macrophage culture and infections

Bone marrow-derived macrophages were isolated from C57BL/6J wild type mice obtained from Jackson Laboratories. The cells were cultured in DMEM supplemented with 10% FBS, 15% L-cell conditioned media, 2 mM L-glutamine, 1 mM sodium pyruvate, and penicillin/streptomycin when needed, and maintained in a 37°C incubator with 5% CO_2_. To induce the formation of foamy macrophages, macrophages were pre-treated with macrophage medium supplemented with oleate/albumin (0.42 mM sodium oleate, 0.35% BSA) complexes 24 hours prior to infection, following a previously established protocol^35,36^. Macrophage infections with Mtb were carried out as previously described^2,58^. For enumeration of colony forming units (CFUs), macrophages were lysed using a solution of water containing 0.01% sodium dodecyl sulfate, and serial dilutions plated on 7H10 agar plates supplemented with 100 μg/ml cycloheximide.

#### Mouse Mtb infections

C3HeB/FeJ wild type mice from Jackson Laboratories were intranasally infected with 10^3^ CFUs of Mtb in a volume of 35 μl, while under light anesthesia with 2% isoflurane^9,11,20^. Following sacrifice using CO_2_ at 2 or 6 weeks post-infection, the left lobe and accessory right lobe of the lungs were homogenized in PBS containing 0.05% Tween 80. Serial dilutions of the homogenates were plated on 7H10 agar plates supplemented with 100 μg/ml cycloheximide for the quantification of CFUs.

#### Confocal immunofluorescence microscopy

Fixed murine lung lobes were processed and stained as previously described^9,61^. DAPI was used at 1:500 (Invitrogen), Bodipy 493/503 was used at 1:500 (Invitrogen), and Alexa Fluor 647 phalloidin was used at 1:50 (Invitrogen). Imaging was performed on a Leica SP8 spectral confocal microscope with 0.5 µm z-steps, and images reconstructed into 3-dimensions with Volocity software (Quorum Technologies).

#### Quantification and statistical analysis

GraphPad Prism was used for all statistical analyses. Details of statistical tests performed are indicated in the figure legends. p<0.05 was considered significant.

## SUPPORTING INFORMATION FIGURE AND TABLE LEGENDS

**Figure S1. PhoPR deletion dampens Mtb response to acidic pH and Cl^−^, in the absence or presence of cholesterol**. Log-phase WT, Δ*phoPR*, and *phoPR*\* (complemented strain) Mtb were exposed for 4 hours to (A) 7H9 or cholesterol media at pH 5.7, or (B) 7H9 or cholesterol media at pH 7 + 250 mM NaCl, along with 7H9, pH 7 as the control condition. Fold change is as compared to the 7H9, pH 7 condition in all cases. *sigA* was used as the control gene, and data are shown as means ± SD from 3 technical replicates, representative of 3 experiments. p-values were obtained with an unpaired t-test with Welch’s correction and Holm-Sidak multiple comparisons, N.S. not significant, * p<0.05, *** p<0.001, **** p<0.0001.

**Table S1. Results of CDC1551(*rv2390c’*::luciferase) transcription factor overexpression pH/K^+^ screen.**

**Table S2. Comparison of effect of acidic pH on genes differentially expressed ≥1 log_2_-fold in cholesterol in WT Mtb, after 4 hours exposure to cholesterol media at pH 5.7 or pH 7.**

**Table S3. Comparison of effect of cholesterol on genes differentially expressed ≥1 log_2_-fold in acidic pH in WT Mtb, after 4 hours exposure to cholesterol, pH 5.7 or 7H9, pH 5.7 media.**

**Table S4. Comparison of effect of *mce3R* deletion on genes differentially expressed ≥1 log_2_-fold in acidic pH in WT Mtb, after 4 hours exposure to 7H9, pH 5.7 medium.**

**Table S5. Comparison of effect of *mce3R* deletion on genes differentially expressed ≥1 log_2_-fold in cholesterol in WT Mtb, after 4 hours exposure to cholesterol, pH 7 medium.**

**Table S6. Comparison of effect of *mce3R* deletion on genes differentially expressed ≥1 log_2_-fold in cholesterol, pH 5.7 in WT Mtb, after 4 hours exposure to cholesterol, pH 5.7 medium.**

