## Supplementary material for "*Mycobacterium tuberculosis* response to cholesterol is integrated with environmental pH and potassium levels via a lipid utilization regulator": Figure S1

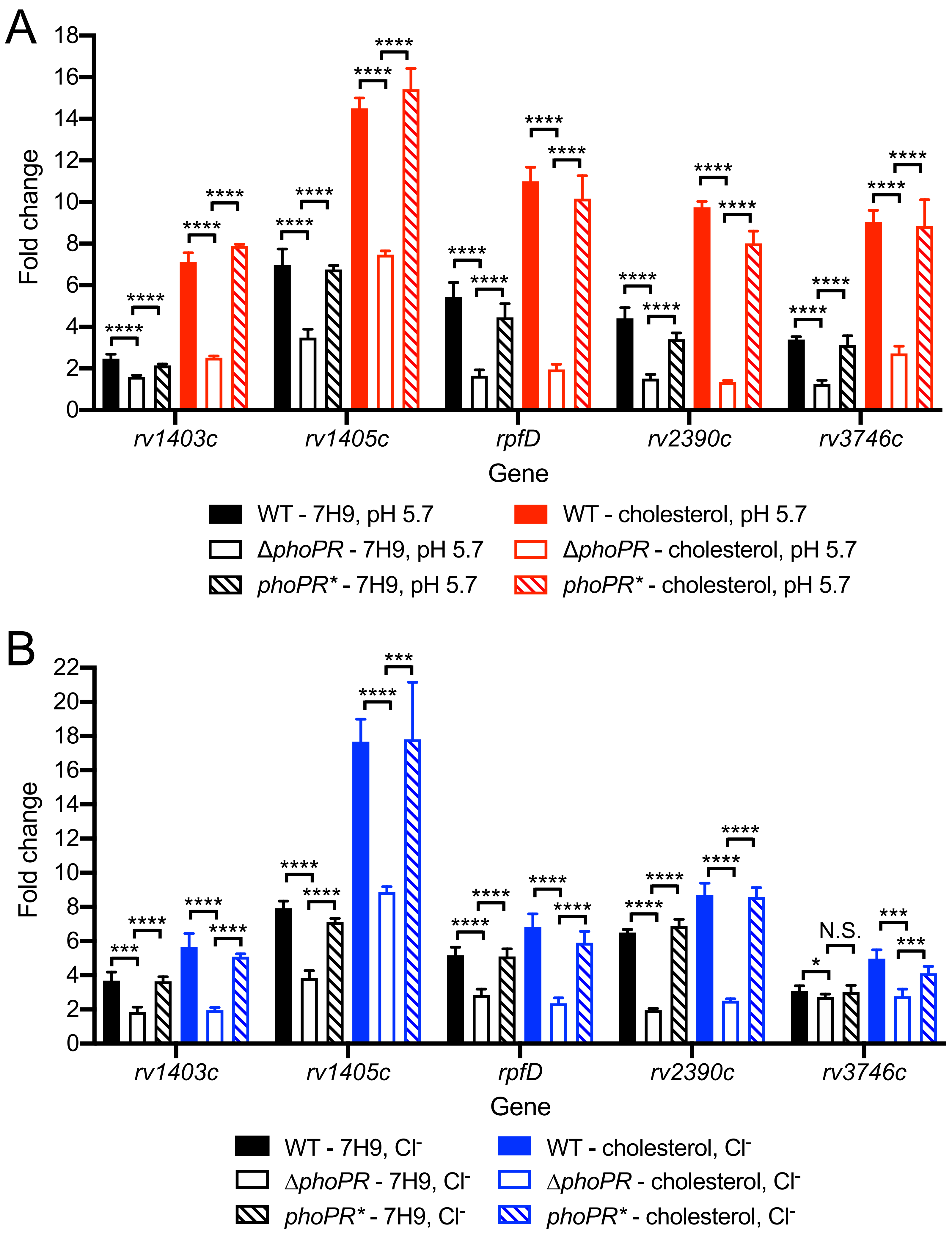


**Figure S1. PhoPR deletion dampens Mtb response to acidic pH and $Cl^-$, in the absence or presence of cholesterol**. Log-phase WT, Δ*phoPR*, and *phoPR** (complemented strain) Mtb were exposed for 4 hours to (A) 7H9 or cholesterol media at pH 5.7, or (B) 7H9 or cholesterol media at pH 7 + 250 mM NaCl, along with 7H9, pH 7 as the control condition. Fold change is as compared to the 7H9, pH 7 condition in all cases. *sigA* was used as the control gene, and data are shown as means ± SD from 3 technical replicates, representative of 3 experiments. p-values were obtained with an unpaired t-test with Welch's correction and Holm-Sidak multiple comparisons, N.S. not significant, * $p<0.05$, *** $p<0.001$, **** $p<0.0001$.
